# Age-Related Patterns of DNA Methylation Changes

**DOI:** 10.1101/2024.12.10.627727

**Authors:** Kevin Yueheng Chen, Wenshu Wang, Hari Naga Sai Kiran Suryadevara, Gang Peng

## Abstract

Epigenetic clocks have achieved significant success in aging research, but they often assume linear methylation changes with age and lack biological interpretability. Using data from 4,641 samples across 23 GEO datasets, we analyzed 1,557 CpGs from nine widely used clocks with minimal overlap, and identified consistent age-associated methylation patterns. We then identified 19,432 age-associated CpGs (aaCpGs) that were strongly correlated with age and showed high consistency between sexes, with faster methylation changes observed in males. Most aaCpGs were identified during early and late life stages, indicating accelerated epigenetic changes during development and aging. No specific genomic enrichment was observed. Clustering analysis revealed four distinct, non-linear age-related methylation trajectories. These findings underscore the complexity of epigenetic aging and suggest that current clocks may overlook important dynamic patterns, particularly after age 65. Incorporating these insights could improve the accuracy and biological relevance of future epigenetic clocks, especially for use across diverse age ranges and populations.

## Background

DNA methylation is a biological process in which methyl groups are attached to DNA molecules, most commonly at CpG dinucleotides, and is functionally linked to gene expression regulation, cellular differentiation, and genome stability^1,2^. Over the past three decades, the relationship between DNA methylation and aging has been extensively explored^3–6^, particularly with the development of DNA methylation clocks designed to estimate biological age^7–14^. Rather than focusing on individual CpGs, these epigenetic clocks are typically constructed from dozens or even hundreds of CpGs selected through feature selection methods such as lasso and elastic net^15,16^. Epigenetic clocks have demonstrated superior accuracy in predicting biological age compared to telomere length, transcriptomic, proteomic, and metabolomic predictors, as well as composite biomarkers^17^. The discrepancy between epigenetic age and chronological age, known as age acceleration, has been associated with mortality and the risk of various diseases^14,18-22^, and it is frequently used as an indicator of individual health status. Studies have shown that a healthy lifestyle, including regular physical activity and a diet rich in lean meats, can mitigate epigenetic age acceleration^23,24^.

Although epigenetic clocks are widely used and perform well in various aging-related studies, their underlying biological mechanisms remain largely unknown^25^. While the epigenetic age estimated by most clocks shows a strong correlation with chronological age^26,27^, there is minimal overlap in the CpGs used across different clocks (**Supplementary Figure 1A**), making it challenging to determine the biological functions of the selected CpGs. Given that epigenetic clocks with distinct sets of CpGs yield similar performance, this suggests there are much more CpGs than the CpGs in the epigenetic clocks that are correlated with age, and these age-related CpGs should also be highly correlated. It is crucial to discover the patterns of DNA methylation changes of the CpGs in these clocks with age to obtain deeper insights into the epigenetic clock and its relationship with aging. While most epigenetic clocks are constructed using linear models, studies have indicated that some DNA methylation changes occur nonlinearly with age, particularly showing accelerated changes in the pediatric age range^28,29^. Moreover, some CpGs included in epigenetic clocks do not exhibit a linear correlation with age (**Supplementary Figure 1B**). Therefore, it is essential to investigate DNA methylation changes at CpG sites that exhibit both linear and nonlinear age-related patterns.

In this study, we aimed to identify patterns of DNA methylation changes associated with age. We downloaded DNA methylation data for 4,641 samples from Gene Expression Omnibus (GEO), analyzed CpGs from nine epigenetic clocks, and identified patterns of age-related methylation changes. We then extended our analysis to additional CpGs and observed four unique patterns across the full sample set, as well as within male and female groups separately. DNA methylation changes occurred more rapidly before the age of 20 and after 60. Additionally, we observed that DNA methylation age progressed faster in males compared to females. These findings demonstrate the strong correlation between DNA methylation and processes of development and aging.

## Results

### Summary of DNA Methylation Data

We collected a total of 4,932 samples from 23 publicly available datasets on the Gene Expression Omnibus (GEO) (**Supplementary Table 1**), applying inclusion criteria of non-cancer status and profiling using either the Illumina 450K or Illumina EPIC arrays. After data harmonization and cleaning, 393,628 CpGs from 4,641 samples (4,489 if only counting those with gender metadata), with ages ranging from 0 to 80 years, were retained for further analysis. (**Figure 1, Supplementary Table 2**). The distribution of ages was relatively the same between male and female samples, but overall, it is not distributed normally; there were significantly more samples between 0 years and 20 years as well as between 40 to 65 years compared to between 20 to 40 years (**Supplementary Figure 2A**). The ethnic composition of the samples was 20.3% White, 3.1% Black, 5.0% Asian, 1.0% Hispanic, and 70.6% Undisclosed (**Supplementary Table 1**). A matrix of DNA methylation level of the 393,638 CpGs (row) from 0 to 80 years (column) old were created (**Figure 1, Details in the Methods**).

**Figure 1.**
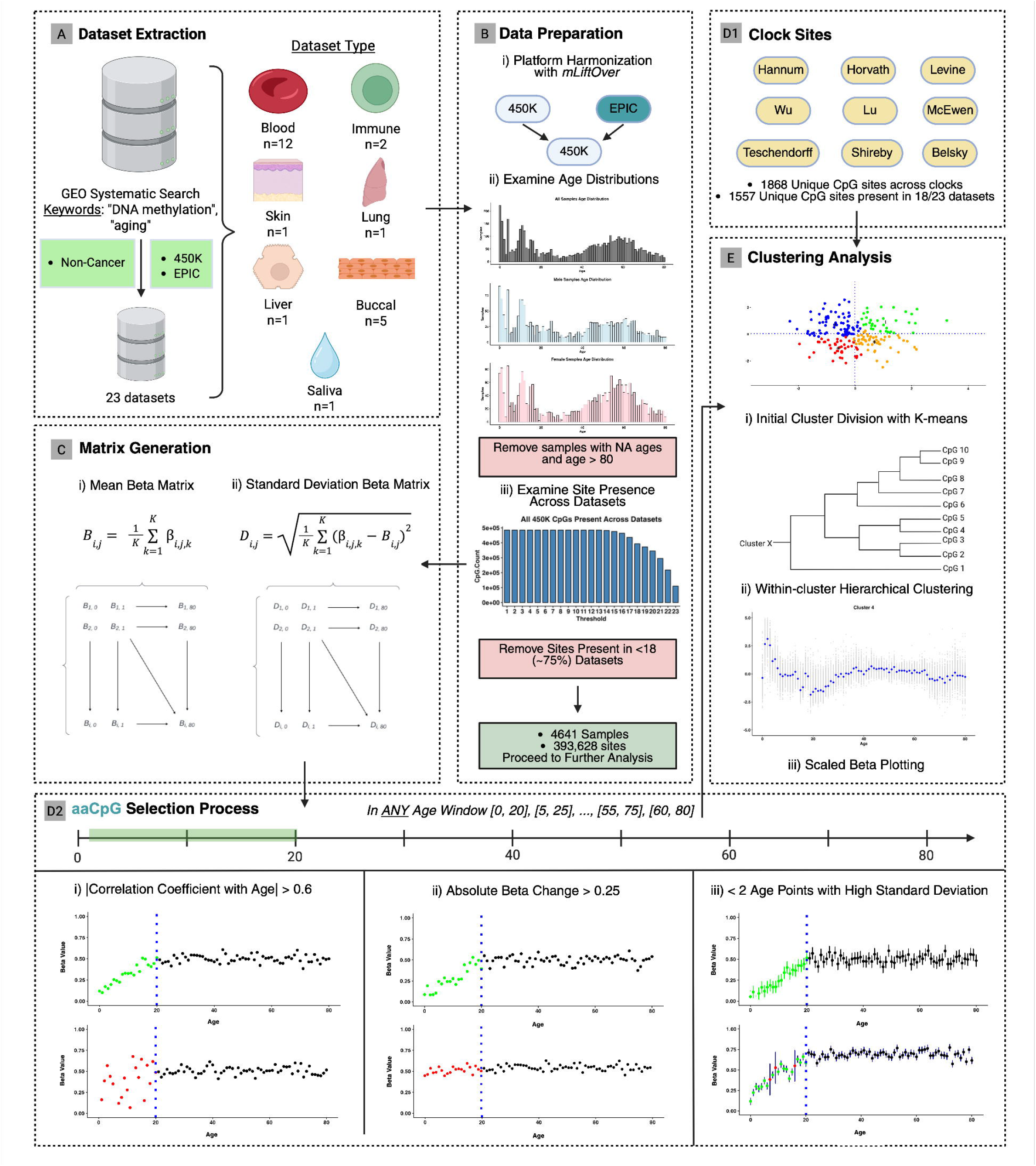
Visual workflow of data analysis. A) DNA methylation data acquisition of our complete cohort. DNA methylation data of 23 datasets from 7 tissues were download from GEO. B) Workflow oWe f data cleaning. C) β value matrix generation. Two different types of reduced matrices were generated: mean and standard deviation of β value. D) Extraction of CpGs for analysis. D1, CpGs from major epigenetic clocks; D2, Custom age associated CpGs (aaCpGs) selection. aaCpGs must fulfill 3 criteria: high correlation coefficient, high absolute beta change, and <2 age points with high standard deviation of beta values. E) K-means clustering of CpGs into groups followed by hierarchical clustering within each group. The changes of DNA methylation with age were illustrated using scatter plots.

### Patterns of CpGs in Epigenetic Clocks Changing with Age

We first investigated the association between age and CpGs from nine highly cited clocks built from different background: Hannum^8^, Horvath^7^, Levine (PhenoAge)^11^, Lu^30^, McEwen (PedBE)^10^, Teschendorff^31^, Shireby^32^, Belsky^13^ (**Supplementary Table 3**). While the training data of most clocks focused on adults, Wu and McEwen’s clocks were included to represent the changes between DNA methylation and age at early development stage. The clock CpGs were filtered if they were not present in >18 (~75%) of the 23 datasets (**Supplementary Figure 2B**). The 1,557 CpGs from the nine clocks after filtering could be clustered into four groups (**Figure 2A**). All CpGs in these four groups exhibited significant changes before the age of 30, with relatively smaller changes observed afterward. In clusters 2-4, minor changes were detected after the age of 65. CpGs in cluster 1 had the most sophisticated changes, decreasing at first then increasing around 10 years old, reaching the peak around 20 and then decreasing till 30. CpGs in cluster 2 were consistently high at first but decreased around 15 years old. Both clusters 3 and 4 displayed a pattern where DNA methylation initially decreased, then increased after age 20 till about 30 years old, while CpGs in cluster 4 had larger variation. We also found large DNA methylation change between age 0 and 1 for cluster 1, 3 and 4 (**Figure 2B**).

**Figure 2.**
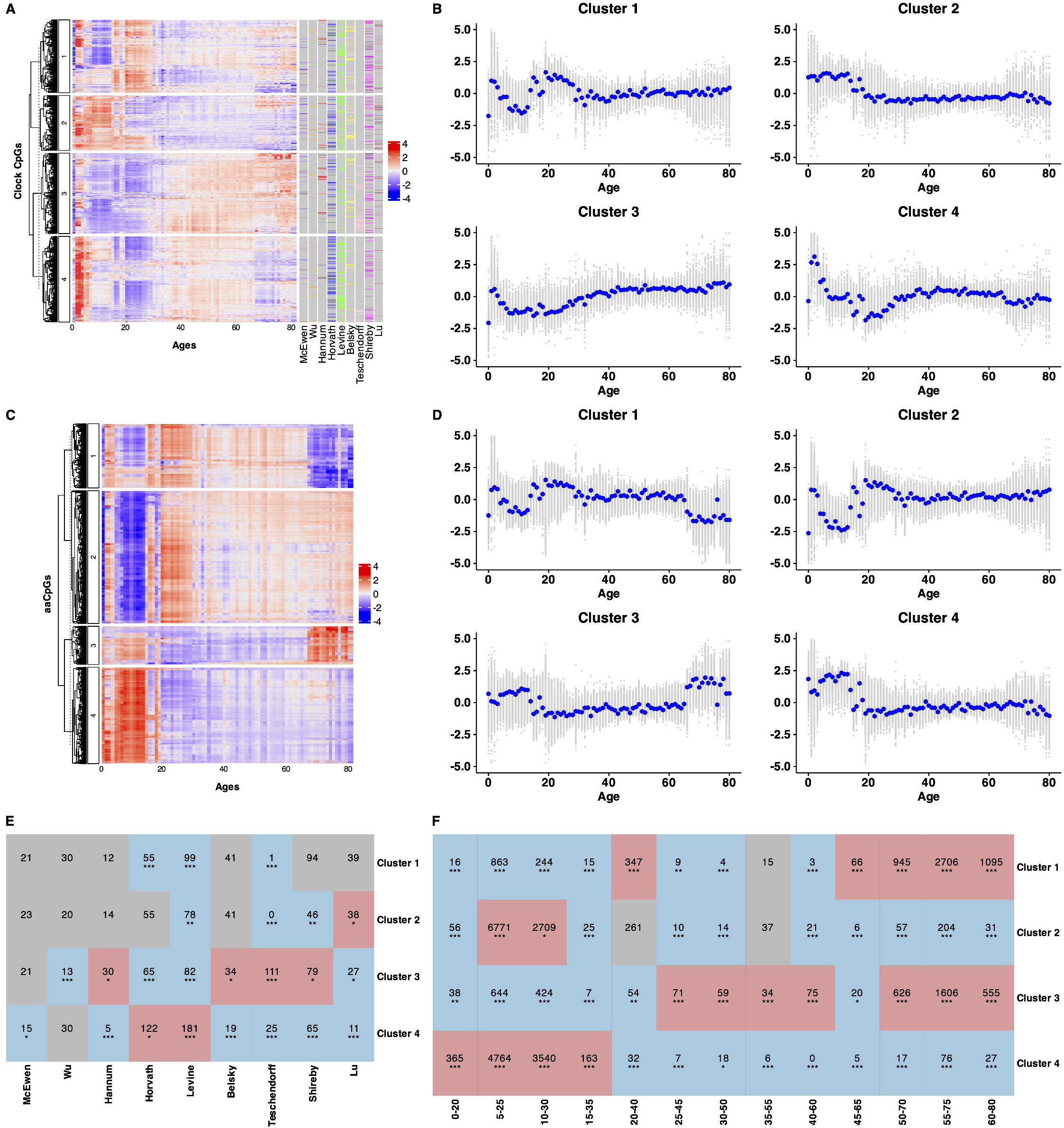
Profiles of CpGs from major epigenetic clocks and aaCpGs. A) Clustering of CpGs from major epigenetic clocks with annotation of origins. B) Scatter plot of the DNA methylation changes of clock CpGs with age within each cluster. C) Clustering of aaCpGs. D) Scatter plot of DNA methylation changes of aaCpG with age within each cluster. E) Number of clock CpGs partitioned in different clusters. F) Number of aaCpGs selected in different age widow within each cluster. Enrichment analysis was performed to assess CpG enrichment for each cluster in both panels E and F. Significant results (Fisher’s exact test with Bonferroni correction, adjusted p-value < 0.05) are shown in red for overrepresentation and blue for underrepresentation.

Although Wu and McEwen’s clocks were built from an adolescent population, we did not find much enrichment or diminishment in any clusters except less CpGs in cluster 3 for Wu’s clock. Belsky and Lu’s clocks were built with participants with age from 26-45 and 53-73 respectively, and more CpGs were found in cluster 2 and less CpGs were found in cluster 4. Most CpGs in Teschendorff’s clock were in cluster 3 with few in cluster 1, 2, and 4 (**Figure 2E**).

### Age-associated CpGs (aaCpGs)

A substantial number of aging-associated CpGs (aaCpGs) are not captured by existing epigenetic clocks. To systematically identify these CpGs while accounting for the nonlinear dynamics of epigenetic aging^33^, we employed a sliding window approach across the age spectrum. Specifically, we divided chronological age into 13 overlapping windows ([0–20], [5– 25], [10–30], …, [55–75], [60–80]) and, within each window, identified CpGs that exhibited strong correlations with age. This window-based strategy allowed for the detection of CpGs associated with age in specific developmental or aging phases, rather than limiting the analysis to CpGs linearly associated with age across the entire lifespan.

We identified aaCpGs across all samples, as well as separately for males and females. The number of aaCpGs detected within each age window is summarized in **Supplementary Table 4**. The majority of aaCpGs were found in windows representing early life (under 40 years, corresponding to development and growth) and later life (over 50 years, corresponding to aging). In contrast, few or no aaCpGs were detected in midlife windows, including [25–45], [30–50], [35–55], [40–60], and [45–65]. The change of DNA methylation were relatively small in the midlife windows (**Supplementary Figure 3**). To enable further analysis of DNA methylation changes within these midlife windows, we selected the top 100 CpGs with the highest correlation coefficients with age in cases where fewer than 100 aaCpGs were identified (**Table 1**). Many aaCpGs detected in each window were window-specific, with overlaps largely occurring between neighboring windows, especially between [5-25] and [10-30]. Interestingly, we also observed a subset of aaCpGs that overlapped between windows associated with early development and later aging, particularly in [5–25], [10–30], [50–70], and [55–75] (**Figure 3A**), suggesting shared regulatory elements across different stages of life. The patterns of overlaps of aaCpGs found in different age window in male-only and female-only were similar to the overall cohort with larger proportional of overlap between [10–30] and [15–35] in female (**Supplementary Figure 4A-B**).

**Table 1.** Number of aaCpGs selected for whole cohort, male-only, and female-only in different age windows.

| Age Window | # aaCpGs |  |  |  |
| --- | --- | --- | --- | --- |
|  | All | Male | Female | Overlap (M/F) |
| [0, 20] | 485 | 1,237 | 498 | 358 |
| [5, 25] | 13,106 | 20,693 | 5,539 | 5,153 |
| [10, 30] | 6,957 | 7,981 | 5,048 | 3,300 |
| [15, 35] | 212 | 100 | 522 | 15 |
| [20, 40] | 702 | 1,339 | 301 | 134 |
| [25, 45] | 100 | 100 | 100 | 0 |
| [30, 50] | 100 | 100 | 100 | 0 |
| [35, 55] | 100 | 100 | 100 | 2 |
| [40, 60] | 100 | 100 | 100 | 3 |
| [45, 65] | 100 | 100 | 100 | 2 |
| [50, 70] | 1,655 | 472 | 978 | 35 |
| [55, 75] | 4,617 | 8,410 | 2,463 | 1643 |
| [60, 80] | 1,718 | 9,662 | 979 | 850 |
| Total | 19,423 | 32,210 | 11,422 | 8105 |

**Figure 3.**
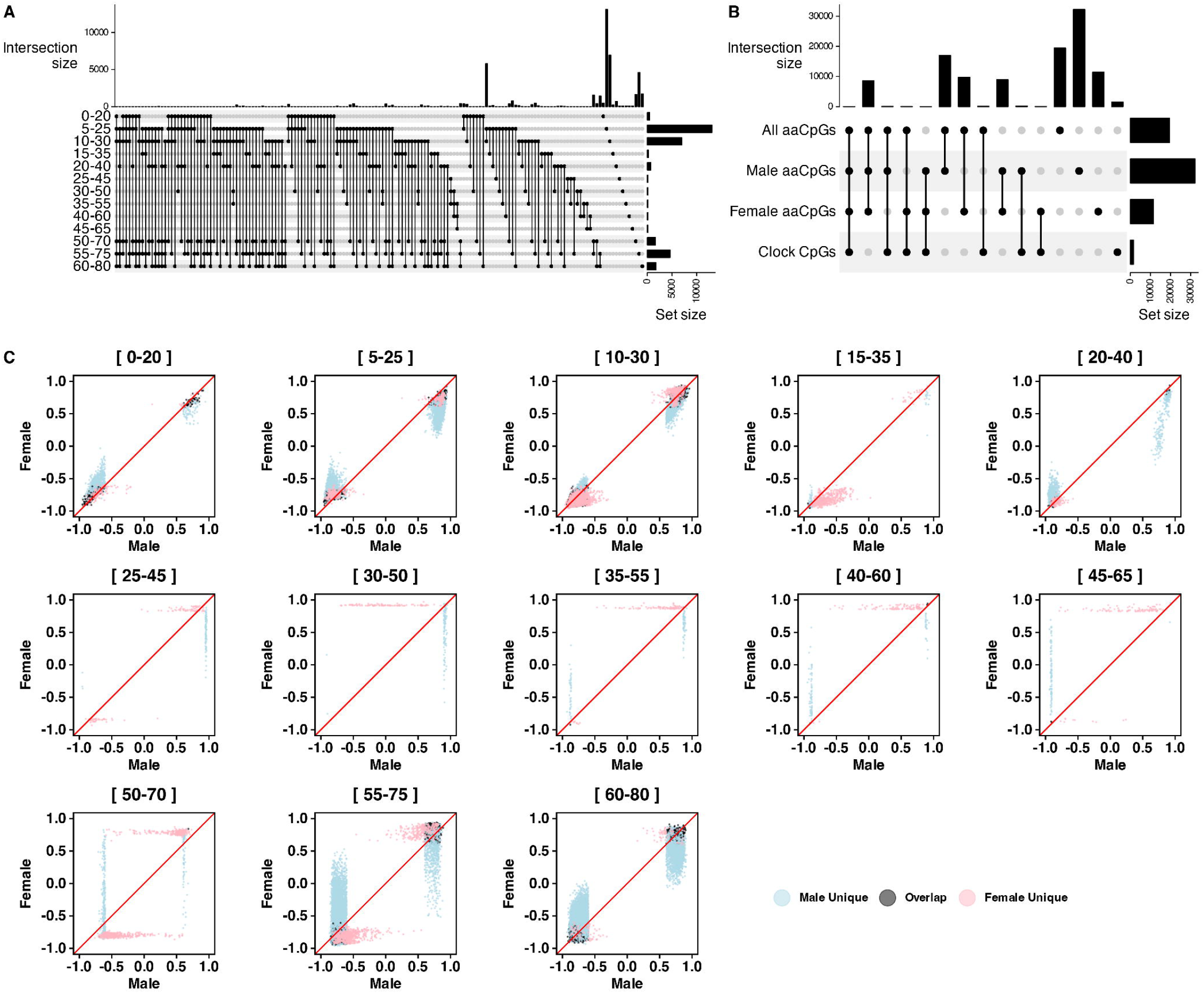
Depictions of aaCpGs selected across different age windows with stratification of sex. A) Overlaps of aaCpGs selected in different age windows. B) Overlap of aaCpGs among the whole cohort, male-only, female-only, and clock CpG sets. C) Consistency of aaCpGs between male and female in different age window. The correlation coefficients between DNA methylation and age within each age window for males and females are shown on the x-axis and y-axis, respectively.

We observed a greater number of aaCpGs in males (n = 32,210) compared to females (n = 11,422), with the most notable difference occurring in the [5–25] age window (**Table 1, Figure 3B**). Most aaCpGs identified in females overlapped with those found in the combined sample or in males, whereas a substantial proportion of aaCpGs identified in males were unique to that group. To further explore sex-specific differences, we examined the correlation coefficients between DNA methylation and age for aaCpGs identified separately in males and females across each age window. The results demonstrated high consistency, particularly during early life (**Figure 3C**). For most aaCpGs uniquely detected in either sex during early life, the direction of correlation with age was the same in the other group, though the effect size was generally larger in males. This suggests that while the age-related methylation patterns are largely shared, the magnitude of association is stronger in males. In the midlife age windows ([25–45] to [45– 65]), aaCpGs were selected based solely on correlation coefficients rather than the original criteria due to the limited number of significant associations. In this range, there was minimal overlap between male and female aaCpGs (**Table 1**), and the sex-specific CpGs exhibited high inter-individual variability in the opposite sex (**Figure 3C**). In contrast, the later life windows showed more overlap and greater consistency in methylation patterns between males and females compared to midlife, though still lower than that observed in early life. We also compared the identified aaCpGs with the CpGs included in existing epigenetic clocks. Only 112 CpGs (7.2%) in the clocks overlapped with aaCpGs (**Figure 3B**).

To examine potential genomic enrichment, we analyzed the distribution of aaCpGs across the genome. Overall, aaCpGs displayed a distribution pattern similar to that of the full set of 28 million CpGs in the genome, with no clear regional enrichment observed—except for a notable concentration in a specific region on chromosome 6 (**Figure 4A**). Interestingly, the 393,628 CpGs used in our study also showed enrichment in this region and shared a similar overall distribution with the aaCpGs. These findings indicate that, compared to the full set of CpGs analyzed, aaCpGs are not strongly enriched in any specific genomic regions. The aaCpGs selected in male and female separately also have a similar distribution as the aaCpGs selected in all samples (**Figure 4B**).

**Figure 4.**
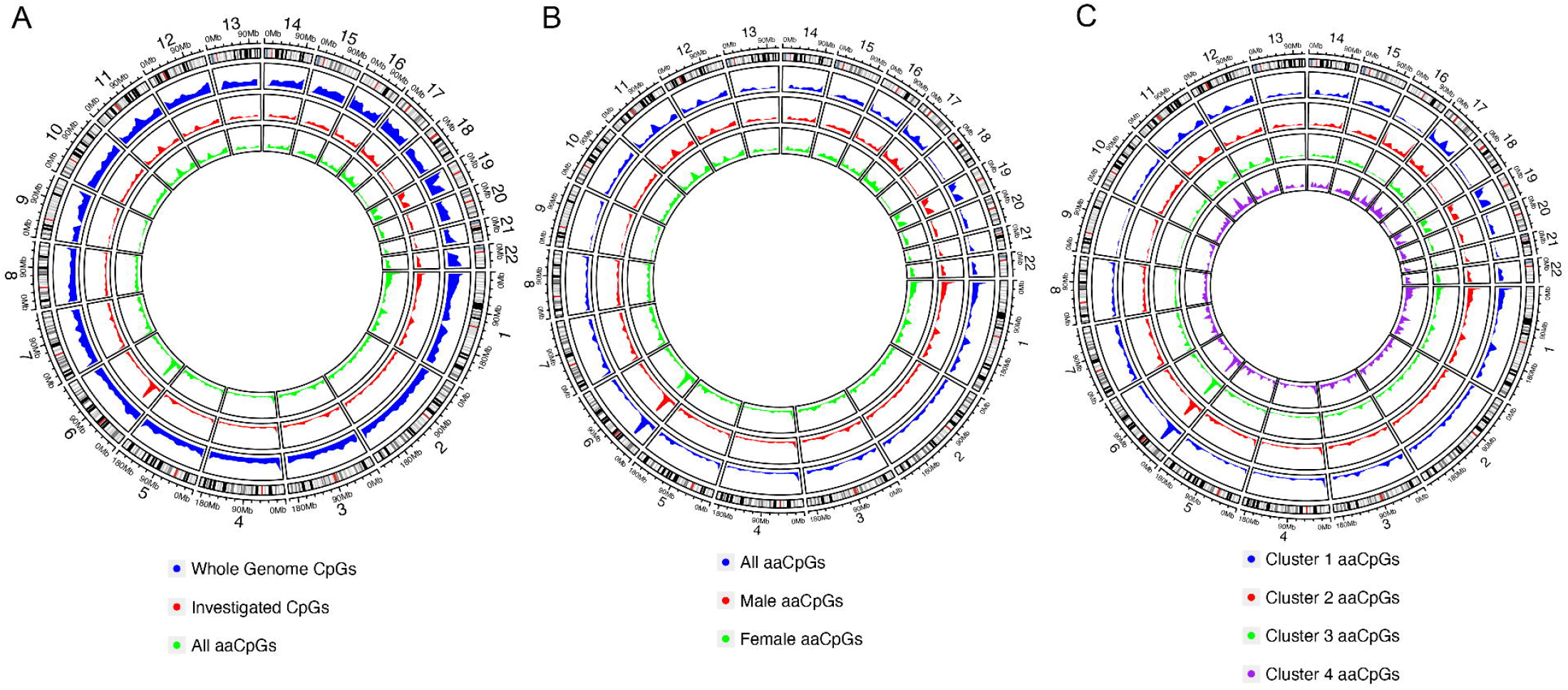
Distribution of aaCpGs in the genome. A) The aaCpGs and the CpGs selected for analysis exhibit similar genomic distributions, both closely resembling the distribution of all CpGs in the genome—except for a notable enrichment in a specific region on chromosome 6. B) The aaCpGs in the whole cohort, male-only, and female-only exhibit similar genome distribution. C) The aaCpGs in 4 clusters resemble similar genome distribution.

### Patterns of Age-Related CpGs Changing with Age

The identified aaCpGs could be clustered into four distinct groups based on their methylation trajectories across age (**Figure 2C**). Notably, cluster 2 and cluster 4 displayed similar age-related patterns to Cluster 1 and Cluster 2 of the clock CpGs, respectively. Clusters 1 and 2 exhibited comparable patterns of methylation change before age 20, although the changes observed in cluster 1 were more modest. A similar relationship was observed between clusters 3 and 4 before age 20. In contrast to Clusters 2 and 4, Clusters 1 and 3 demonstrated more pronounced DNA methylation changes around age 65, suggesting a stronger association with late-life aging processes. Large DNA methylation changes were also found between age 0 and 1, especially in cluster 1 and 2 (**Figure 2D**). Enrichment analysis revealed that aaCpGs identified in windows prior to age 30 were predominantly associated with Clusters 2 and 4, while those identified in windows beyond age 50 were enriched in Clusters 1 and 3 (**Figure 2F**). We further examined the genomic distribution of aaCpGs within each cluster. The distribution patterns were consistent with those observed for the full set of aaCpGs, with no evidence of cluster-specific regional enrichment (**Figure 4C**).

We applied the same clustering analysis to male- and female-specific aaCpGs. The age-associated methylation patterns identified in the overall cohort were largely preserved within each sex (**Figure 5A-D**). In males, Clusters 1 and 3 exhibited marked changes in DNA methylation around age 65. In contrast, the corresponding changes in females were more gradual across the similar age range and did not show a distinct shift around age 52, the typical age of menopause (https://www.nia.nih.gov/health/menopause/what-menopause). This suggests potential sex-specific differences in the magnitude of epigenetic aging dynamics.

**Figure 5.**
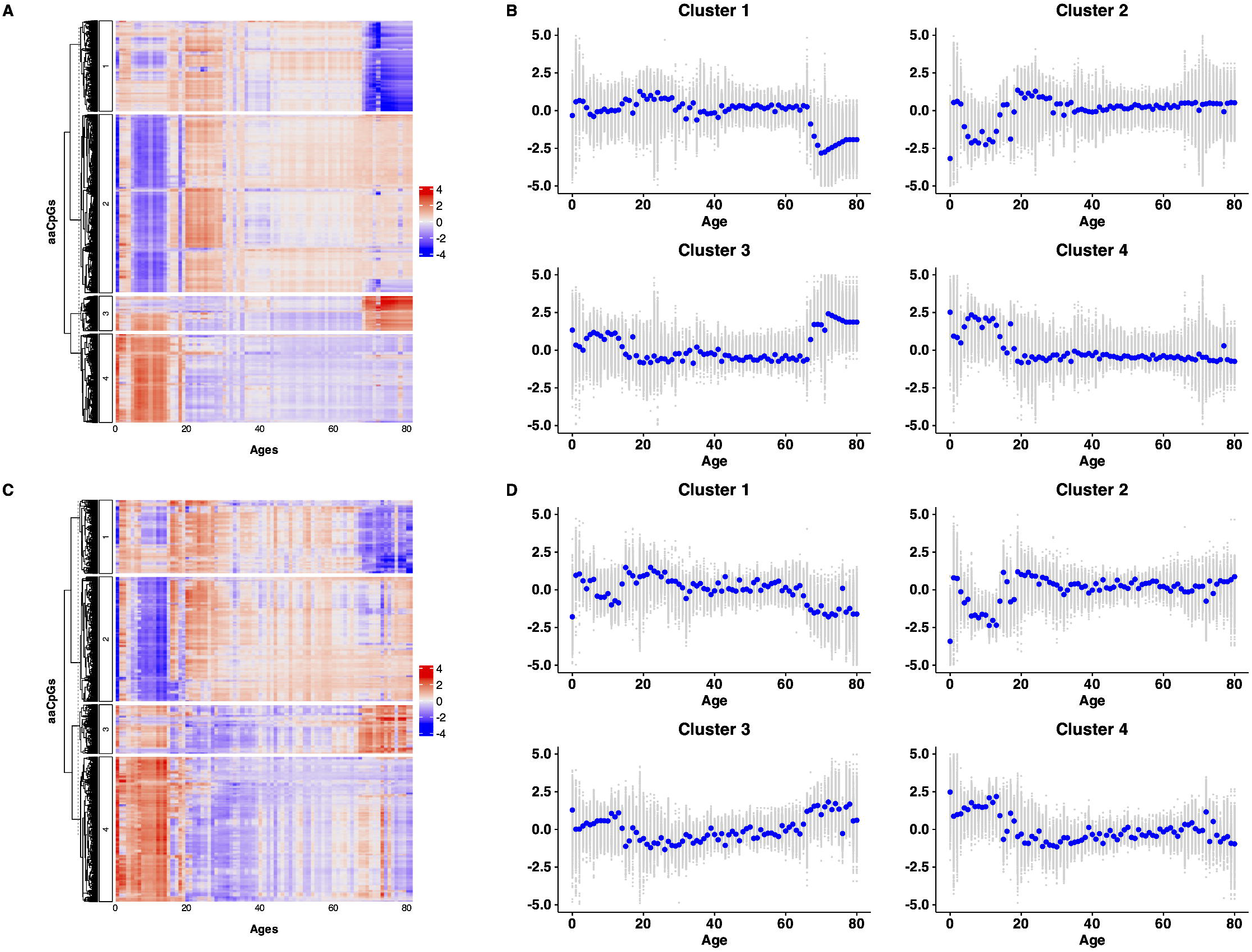
Profile of DNA methylation changes with age of male-only and female only aaCpGs. A) Clustering of male-only aaCpGs. B) Scatter plots showing DNA methylation change of male-only aaCpG with age within each cluster. C) Clustering of female-only aaCpGs. D) Scatter plots showing DNA methylation change of female-only aaCpG with age within each cluster.

## Discussion

The CpGs used across various epigenetic clocks differ substantially (**Supplementary Figure 1A**). In this study, we first examined the shared patterns of age-related DNA methylation changes among CpGs used in epigenetic clocks. Although most of the epigenetic clocks evaluated here were not developed with a focus on young populations, many of the CpGs they include—such as those in the clocks by Belsky and Lu, which were constructed using adults from narrow age ranges (26–45 and 53–73)—exhibit pronounced methylation changes before age 30 and relatively small changes thereafter (**Figure 2A–B**). This trend may help explain the similarity in patterns observed in CpGs from the clocks by Wu and McEwen that focusing on young population (**Figure 2E**). Among the clocks analyzed, Teschendorff’s clock is unique in that most of its CpGs are enriched within a single cluster. This is likely since CpGs in this clock were selected from promoter regions marked by the PRC2 complex and are known to increase with mitotic age^31,34^. Notably, CpGs in cluster 3 show low methylation levels at birth followed by a steady increase over time, aligning with Teschendorff’s selection criteria, except a transient deviation was observed between ages 1 and 10, where methylation levels initially rise sharply between birth and age 1, then decline gradually until around age 10. Teschendorff’s clock was trained without samples less than 19 years old, which may explain the missing capture of the changes.

Then we identified 19,423 CpGs associated with age across all samples with majority found in early and late life. When samples were divided into male and female groups, more age-associated CpGs (aaCpGs) were detected in males than in females (**Table 1**). Although a small proportion of aaCpGs overlapped between the two groups (**Table 1, Figure 3B**), most aaCpGs selected within each age window exhibited the same direction of correlation with age for both males and females (**Figure 3C**). The correlation between DNA methylation and age was generally weaker in females than in males, explaining the higher number of aaCpGs observed in males. This may also account for the faster epigenetic age acceleration seen in males compared to females^8,26^. In general, the aaCpGs detected in male and female were consistent with each other. This consistency supported the validity of our findings. While males and females exhibit similar patterns of DNA methylation change with age, the rate of change is slower in females.

There was limited overlap between aaCpGs and CpGs used in established epigenetic clocks (**Figure 3B**). This discrepancy may be due to the inclusion of some clock CpGs that exhibit weaker correlations with age or greater variability in methylation levels (**Supplementary Figure 1B**). In contrast, aaCpGs were identified based on their strong associations with age within specific 20-year age windows. Consequently, CpGs that display consistent but modest changes in methylation across the lifespan may not meet the selection criteria for aaCpGs, yet they could still be included in epigenetic clocks due to their cumulative association with chronological age. It is a limitation of the current study that does not include CpGs with cumulative association with chronological age.

The genomic distribution of aaCpGs closely mirrors that of CpGs across the entire genome, except for a notable enrichment on chromosome 6p (**Figure 4A**). This enrichment arises from the fact that aaCpGs were selected from a filtered set of 393,628 CpGs, which itself reflects the probe design bias of the Illumina 450K array. Similar genomic distributions were observed for aaCpGs identified in the male-only and female-only groups, as well as across the four aaCpG clusters (**Figure 4B–C**). Overall, these findings suggest that aaCpGs are broadly distributed across the genome, with no significant regional enrichment. On average, approximately one in every 20 CpGs (all samples: 393,628/19,423 ≈ 20; males: 393,628/32,210 ≈ 12; females: 393,628/11,422 ≈ 34) exhibits a strong age association. This indicates that age-related DNA methylation changes are widespread rather than confined to specific genomic regions. A limitation of this analysis is that it is based on a preselected subset of CpGs. Further validation using whole-genome methylation approaches, such as whole-genome bisulfite sequencing or Oxford Nanopore sequencing, is needed to confirm these findings and rule out array-based biases.

Four distinct clusters of aaCpGs were identified (**Figure 2C–D**). All four clusters exhibited substantial DNA methylation changes before the age of 20, while only two showed marked changes around age 65. In contrast, methylation changes during midlife were relatively modest (**Supplementary Figure 3, Supplementary Table 1**). Similar age-associated patterns were observed in the male-only and female-only analyses, although females displayed smoother transitions in late life compared to males (**Figure 3**). These findings underscore the nonlinearity of age-related DNA methylation changes, diverging from the linear assumptions made in most epigenetic clocks and aligning with recent multi-omics studies that reported nonlinear aging trajectories^33^. Interestingly, while previous studies reported significant molecular changes around ages 44 and 60^33^, our results revealed a pronounced shift only around age 65 in adults. In early life, we also observed particularly large methylation changes between birth and one year of age for some aaCpGs. Notably, we did not detect strong methylation shifts around age 50 in females, which may be attributed to increased variability in CpGs associated with menopause; such variation likely led to their exclusion during aaCpG selection for the corresponding age windows.

Compared to the CpGs used in existing epigenetic clocks, two novel clusters of aaCpGs were identified that exhibited substantial DNA methylation changes after the age of 65 (**Figure 2C–D**). While these two late-life clusters also showed some methylation changes during early life, the magnitude of those changes was smaller than that observed in the other two clusters, which were primarily associated with early life. Thus, the aaCpGs can be broadly categorized into two groups: those linked primarily to early-life methylation dynamics and those showing changes in both early and late life. The latter group may explain the observed overlap of aaCpGs between younger (5–30 years) and older (50–75 years) age windows (**Figure 3A**). In contrast, the CpGs included in existing epigenetic clocks showed limited methylation change in late life (**Figure 2A–B**), with minimal overlap with the aaCpGs (**Figure 3B**). This discrepancy may be due to limitations in the training datasets, which often contain fewer samples over age 70 (as seen in our data, **Supplementary Figure 2**). This suggests that current epigenetic clocks may fail to capture critical late-life epigenetic alterations, reducing their sensitivity in older populations. Indeed, for most individuals aged 65 and above, epigenetic age estimates derived from existing clocks were lower than chronological age and showed only modest variation beyond age 50 (**Supplementary Figure 5**), which limits their utility for aging research in elderly cohorts. It is essential to incorporate CpGs that exhibit substantial changes in late life into epigenetic clocks, or alternatively, to develop a new clock specifically tailored for the elderly population.

In summary, we identified aaCpGs that exhibit strong correlations with chronological age, with a greater number detected in males—potentially reflecting faster epigenetic aging in this group. Although there was limited overlap in aaCpGs between sexes, the consistency in age-related methylation patterns supports the robustness of our findings. These aaCpGs were broadly distributed across the genome without clear regional enrichment. Clustering analysis revealed distinct, non-linear methylation trajectories: all aaCpGs underwent substantial changes before age 20, while others exhibited notable shifts both in early life and around age 65. These results suggest that age-related methylation changes are widespread and occur at specific developmental and aging stages. Our findings may deepen understanding of the dynamic relationship between DNA methylation and biological aging and inform the development of more accurate and biologically meaningful epigenetic clocks.

## Methods

### Data Availability

All datasets used in this study were publicly available on the Gene Expression Omnibus (GEO). Dataset accessions: GSE32148, GSE36054, GSE40279, GSE50660, GSE50759, GSE51057, GSE53740, GSE61256, GSE67705, GSE73103, GSE80261, GSE85568, GSE89253, GSE90124, GSE94734, GSE106648, GSE114134, GSE124366, GSE137495, GSE138279, GSE62924, GSE51180, GSE30870 (**Supplementary Table 1**). We selected them considering the following criteria: array type (The Illumina HumanMethylation450 BeadChip and HumanMethylationEPIC BeadChip), age metadata present, disease status (exclude samples with cancer), and dataset already processed into beta or M value.

### Data Extraction and Cleaning

Methylation data and patient metadata were systematically extracted and organized using Python libraries GEOparse, pandas, gzip, and math from the Gene Expression Omnibus (GEO).

Platform differences between the EPIC and legacy 450k datasets were corrected for using the mLiftOver package. GSE89253 and GSE50759 were converted from M-values to -values using the equation 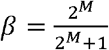.

We removed samples that did not have age metadata and only investigated those between age 0 and age 80. We only investigated CpGs that were present in at least 18/23 (~75%) datasets.

### β Value Matrix Generation

All further analysis was conducted using a condensed version of the datasets. For reference, *B* is a matrix of methylation level of CpGs at different ages, where B_i, j_ is the β value of CpG *i* at age *j*, j ∈ 0, 1, 2, 3, ⋯, 79, 80. If there were multiple samples including CpG *i* at age *j*, we used mean β value, 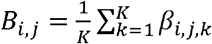, assuming there are *K* samples at age *j* who have data for CpG *i. βi,j,k* is the beta value for CpG *i* at age *j* for sample *k. D* is a variance matrix, where D_i, j_ is the standard deviation of beta values of CpG *i* at age *j*, j ∈ 0, 1, 2, 3, ⋯, 79, 80, 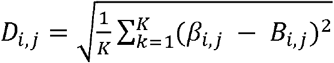. Besides the matrices for the whole cohort, we also generated matrices for male and female separately: *B*_*M*_ (mean beta matrix, only male samples included), *D*_*M*_ (variance beta matrix, only male samples included), *B*_*F*_ (mean beta matrix, only female samples included), *D*_*F*_ (variance beta matrix, only female samples included). We linearly interpolated missing values (~0.5%) in the final matrices.

### Epigenetic Clocks

We investigated CpG sites used across highly cited clocks: McEwen (PedBE)^10^, Wu^35^, Hannum^8^, Horvath (Multi-Tissue Age Estimator)^7^, Levine (PhenoAge)^11^, Belsky (DunedinPACE)^13^, Teschendorff (EpiTOC2)^31^, Shireby (Cortical Clock)^32^, and Lu (DNAmTL)^30^. CpGs used in each clock were extracted from the corresponding paper’s supplementary materials.

### Age-associated CpG sites Selection

We selected a custom set of age-associated CpG sites (aaCpGs) across all datasets using a sliding window approach. A complete set of aaCpGs is defined as all unique CpG sites selected across all age windows. The chronological age from 0 to 80 were divided into the following sliding windows: [0, 20], [5, 25], [10, 30], …, [55, 75], [60, 80]. CpG sites passed our filtering criterion (Figure 1D2) when having (i) |correlation coefficient between β value and age| > 0.6 AND (ii) absolute mean beta change > 0.25 AND (iii) large variation at less than 2 age points in any sliding window. Large variation at CpG site *i* and age *j* can be defined as having standard deviation D_i,j_ larger than 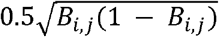, where B_i, j_ is the mean beta value. If the number of aaCpGs for an age window was less than 100, a secondary method was employed. aaCpGs had to fulfill the variation criteria (iii above), the top 100 sites by correlation coefficient were taken, then the top 100 sites by absolute beta change were selected. Different sets of aaCpGs were taken for *B, B*_*M*_, *and B*_*F*_.

### CpG Clustering

Clustering was performed using an integrated k-means and hierarchical clustering approach from R ComplexHeatmap 2.20.0. Default parameters were used with Euclidean distance and complete linkage. Beta values were scaled to ensure age-related patterns were preserved while disregarding their original methylation ranges. The data was initially partitioned into *k* clusters using the k-means algorithm. Hierarchical clustering was conducted independently within each cluster with the clusters then being agglomerated. The optimal number of clusters, k, was determined through a combination of elbow method and an iterative process. Elbow method indicated k ranges from 4 to 7. Beginning with k = 4 clusters, we incrementally increased k by one and evaluated the resulting cluster configurations. If the additional cluster revealed a distinguishable pattern, we proceeded by setting k = k + 1. This process was continued until no further discernible patterns emerged, at which point the iteration was terminated. Clusters were visualized by heatmap and scatter plot of scaled methylation level of CpGs in each cluster at different.

### Enrichment Analysis

We used Fisher’s exact test to assess i) the enrichment of clock CpGs in each cluster and ii) the enrichment of aaCpGs selected in each age window in each cluster. P-values were adjusted using the Benjamini-Hochberg method to identify clusters with significant feature enrichment.

### Genomic Distribution

The circos plot showing the distribution of CpGs was generated using R karyoploteR 1.30.0, with CpGs annotated to their genomic coordinates with R GenomicRanges 1.56.2.

## Supporting information

All Supplementary Figures and Descriptions

All Supplementary Tables

## Code Availability

All code is hosted on our github: https://github.com/PengBioinformaticsLab/AgePatternDNAMethylation

## References

1 Moore, L. D., Le, T. & Fan, G. DNA methylation and its basic function. Neuropsychopharmacology 38, 23–38 (2013).

2 Mattei, A. L., Bailly, N. & Meissner, A. DNA methylation: a historical perspective. Trends Genet 38, 676–707 (2022). 10.1016/j.tig.2022.03.010

3 Fraga, M. F. et al. Epigenetic differences arise during the lifetime of monozygotic twins. Proc Natl Acad Sci U S A 102, 10604–10609 (2005). 10.1073/pnas.0500398102

4 Bollati, V. et al. Decline in genomic DNA methylation through aging in a cohort of elderly subjects. Mech Ageing Dev 130, 234–239 (2009). 10.1016/j.mad.2008.12.003

5 Bell, J. T. et al. Epigenome-wide scans identify differentially methylated regions for age and age-related phenotypes in a healthy ageing population. PLoS Genet 8, e1002629 (2012). 10.1371/journal.pgen.1002629

6 McCartney, D. L. et al. An epigenome-wide association study of sex-specific chronological ageing. Genome Med 12, 1 (2019). 10.1186/s13073-019-0693-z

7 Horvath, S. DNA methylation age of human tissues and cell types. Genome Biol 14, R115 (2013). 10.1186/gb-2013-14-10-r115

8 Hannum, G. et al. Genome-wide methylation profiles reveal quantitative views of human aging rates. Mol Cell 49, 359–367 (2013). 10.1016/j.molcel.2012.10.016

9 Lu, A. T. et al. Universal DNA methylation age across mammalian tissues. Nat Aging 3, 1144–1166 (2023). 10.1038/s43587-023-00462-6

10 McEwen, L. M. et al. The PedBE clock accurately estimates DNA methylation age in pediatric buccal cells. Proc Natl Acad Sci U S A 117, 23329–23335 (2020). 10.1073/pnas.1820843116

11 Levine, M. E. et al. An epigenetic biomarker of aging for lifespan and healthspan. Aging (Albany NY) 10, 573–591 (2018). 10.18632/aging.101414

12 Knight, A. K. et al. An epigenetic clock for gestational age at birth based on blood methylation data. Genome Biol 17, 206 (2016). 10.1186/s13059-016-1068-z

13 Belsky, D. W. et al. DunedinPACE, a DNA methylation biomarker of the pace of aging. Elife 11 (2022). 10.7554/eLife.73420

14 Lu, A. T. et al. DNA methylation GrimAge strongly predicts lifespan and healthspan. Aging (Albany NY) 11, 303–327 (2019). 10.18632/aging.101684

15 Tibshirani, R. Regression shrinkage and selection via the lasso. Journal of the Royal Statistical Society: Series B (Methodological) 58, 267–288 (1996).

16 Zou, H. & Hastie, T. Regularization and variable selection via the elastic net. Journal of the Royal Statistical Society Series B: Statistical Methodology 67, 301–320 (2005).

17 Jylhävä, J., Pedersen, N. L. & Hägg, S. Biological Age Predictors. EBioMedicine 21, 29–36 (2017). 10.1016/j.ebiom.2017.03.046

18 Marioni, R. E. et al. DNA methylation age of blood predicts all-cause mortality in later life. Genome Biol 16, 25 (2015). 10.1186/s13059-015-0584-6

19 Xiao, C. et al. Association of Epigenetic Age Acceleration With Risk Factors, Survival, and Quality of Life in Patients With Head and Neck Cancer. Int J Radiat Oncol Biol Phys 111, 157–167 (2021). 10.1016/j.ijrobp.2021.04.002

20 Jain, P. et al. Analysis of Epigenetic Age Acceleration and Healthy Longevity Among Older US Women. JAMA Netw Open 5, e2223285 (2022). 10.1001/jamanetworkopen.2022.23285

21 Faul, J. D. et al. Epigenetic-based age acceleration in a representative sample of older Americans: Associations with aging-related morbidity and mortality. Proc Natl Acad Sci U S A 120, e2215840120 (2023). 10.1073/pnas.2215840120

22 Fransquet, P. D., Wrigglesworth, J., Woods, R. L., Ernst, M. E. & Ryan, J. The epigenetic clock as a predictor of disease and mortality risk: a systematic review and meta-analysis. Clin Epigenetics 11, 62 (2019). 10.1186/s13148-019-0656-7

23 Quach, A. et al. Epigenetic clock analysis of diet, exercise, education, and lifestyle factors. Aging (Albany NY) 9, 419–446 (2017). 10.18632/aging.101168

24 Fitzgerald, K. N. et al. Potential reversal of epigenetic age using a diet and lifestyle intervention: a pilot randomized clinical trial. Aging (Albany NY) 13, 9419–9432 (2021). 10.18632/aging.202913

25 Kabacik, S. et al. The relationship between epigenetic age and the hallmarks of ageing in human cells. Nature Aging, 1-10 (2022).

26 Crimmins, E. M., Thyagarajan, B., Levine, M. E., Weir, D. R. & Faul, J. Associations of Age, Sex, Race/Ethnicity, and Education With 13 Epigenetic Clocks in a Nationally Representative U.S. Sample: The Health and Retirement Study. J Gerontol A Biol Sci Med Sci 76, 1117–1123 (2021). 10.1093/gerona/glab016

27 Liu, Z. et al. Underlying features of epigenetic aging clocks in vivo and in vitro. Aging Cell 19, e13229 (2020). 10.1111/acel.13229

28 Vershinina, O., Bacalini, M. G., Zaikin, A., Franceschi, C. & Ivanchenko, M. Disentangling age-dependent DNA methylation: deterministic, stochastic, and nonlinear. Sci Rep 11, 9201 (2021). 10.1038/s41598-021-88504-0

29 Alisch, R. S. et al. Age-associated DNA methylation in pediatric populations. Genome Res 22, 623–632 (2012). 10.1101/gr.125187.111

30 Lu, A. T. et al. DNA methylation-based estimator of telomere length. Aging (Albany NY) 11, 5895–5923 (2019). 10.18632/aging.102173

31 Teschendorff, A.E. A comparison of epigenetic mitotic-like clocks for cancer risk prediction. Genome Med 12, 56 (2020). 10.1186/s13073-020-00752-3

32 Shireby, G. L. et al. Recalibrating the epigenetic clock: implications for assessing biological age in the human cortex. Brain 143, 3763–3775 (2020). 10.1093/brain/awaa334

33 Shen, X. et al. Nonlinear dynamics of multi-omics profiles during human aging. Nat Aging (2024). 10.1038/s43587-024-00692-2

34 Yang, Z. et al. Correlation of an epigenetic mitotic clock with cancer risk. Genome Biol 17, 205 (2016). 10.1186/s13059-016-1064-3

35 Wu, X. et al. DNA methylation profile is a quantitative measure of biological aging in children. Aging (Albany NY) 11, 10031–10051 (2019). 10.18632/aging.102399

