## Supplementary material for "Age-Related Patterns of DNA Methylation Changes": All Supplementary Figures and Descriptions

**Supplementary Materials**

**Supplementary Figures**

**Supplementary Figure 1.** A) Overlaps of CpGs in major epigenetic clocks. dB) Association between age and DNA methylation changes at CpGs shared across multiple epigenetic clocks. (cg19722847: Wu, Hannum, Horvath, Levine; cg16867657: McEwen, Hannum, Shireby, Lu; cg09809672: Wu, Hannum, Horvath, Levine; cg06144905: McEwen, Horvath, Levine, Shireby)

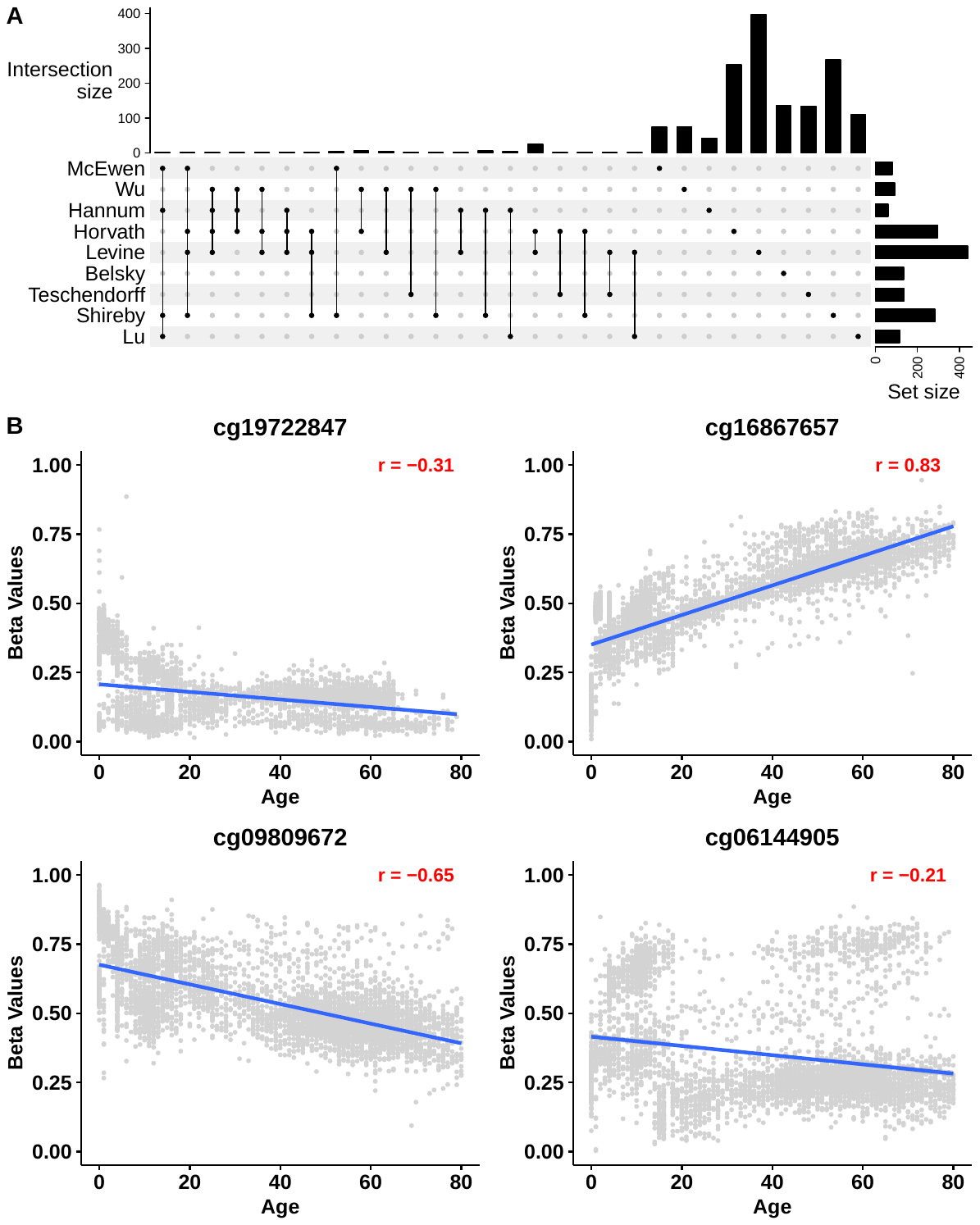

**Supplementary Figure 2.** A) Age distribution of samples in whole cohort, male-only, and female-only. B) Presents of clock CpGs and all CpGs (450K array) we collected from GEO across all 23 datasets. C) Proportion of samples from different datasets at age. Samples from different sources (datasets) could improve the robustness of our findings.

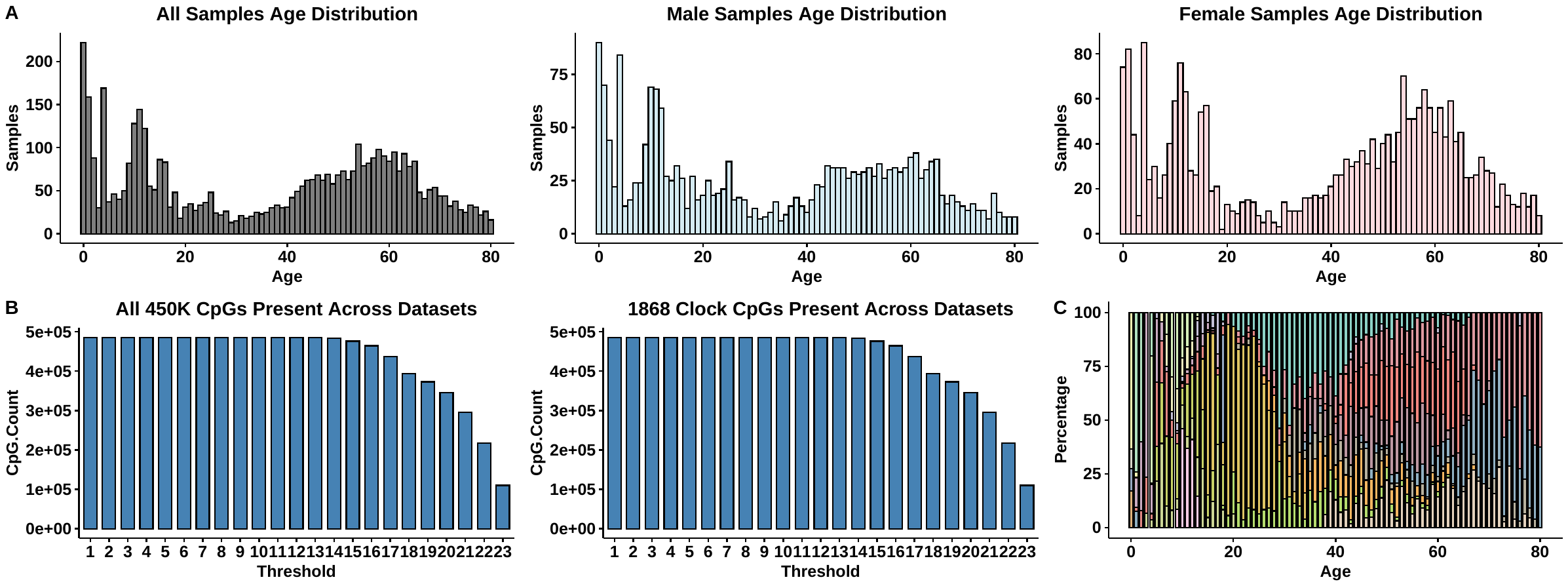

**Supplementary Figure 3**

Absolute beta value change and correlation coefficient between beta value and age for all CpGs in different age windows of all sample, male-only, and female-only.

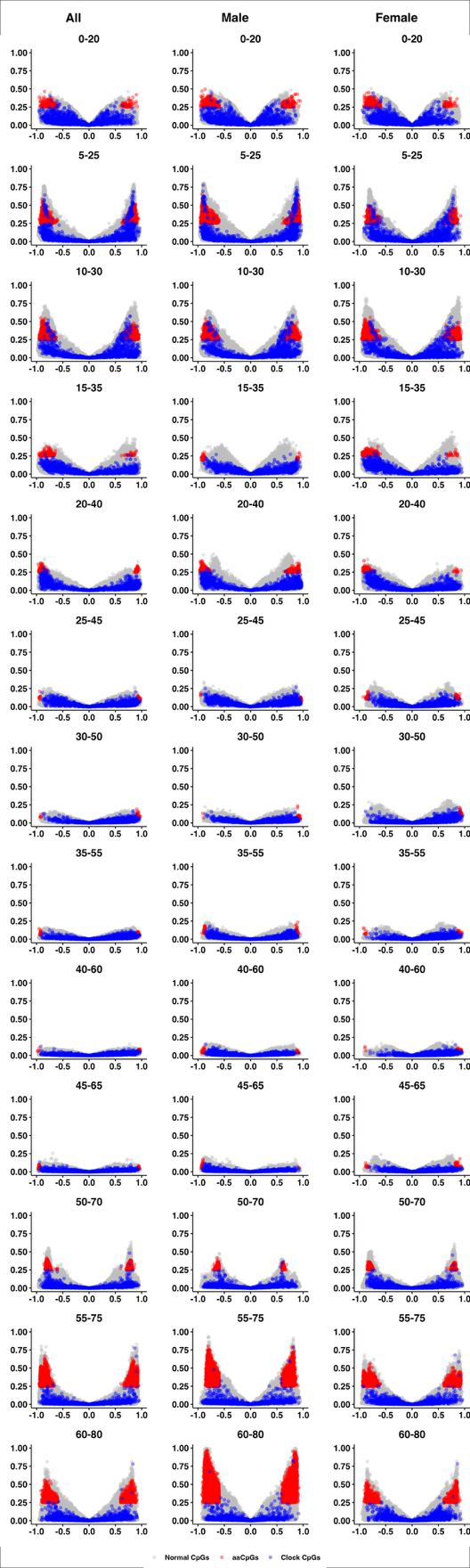

**Supplementary Figure 4.**

Overlap of aaCpGs selected in different age windows. A) Male-only. B) Female-only.

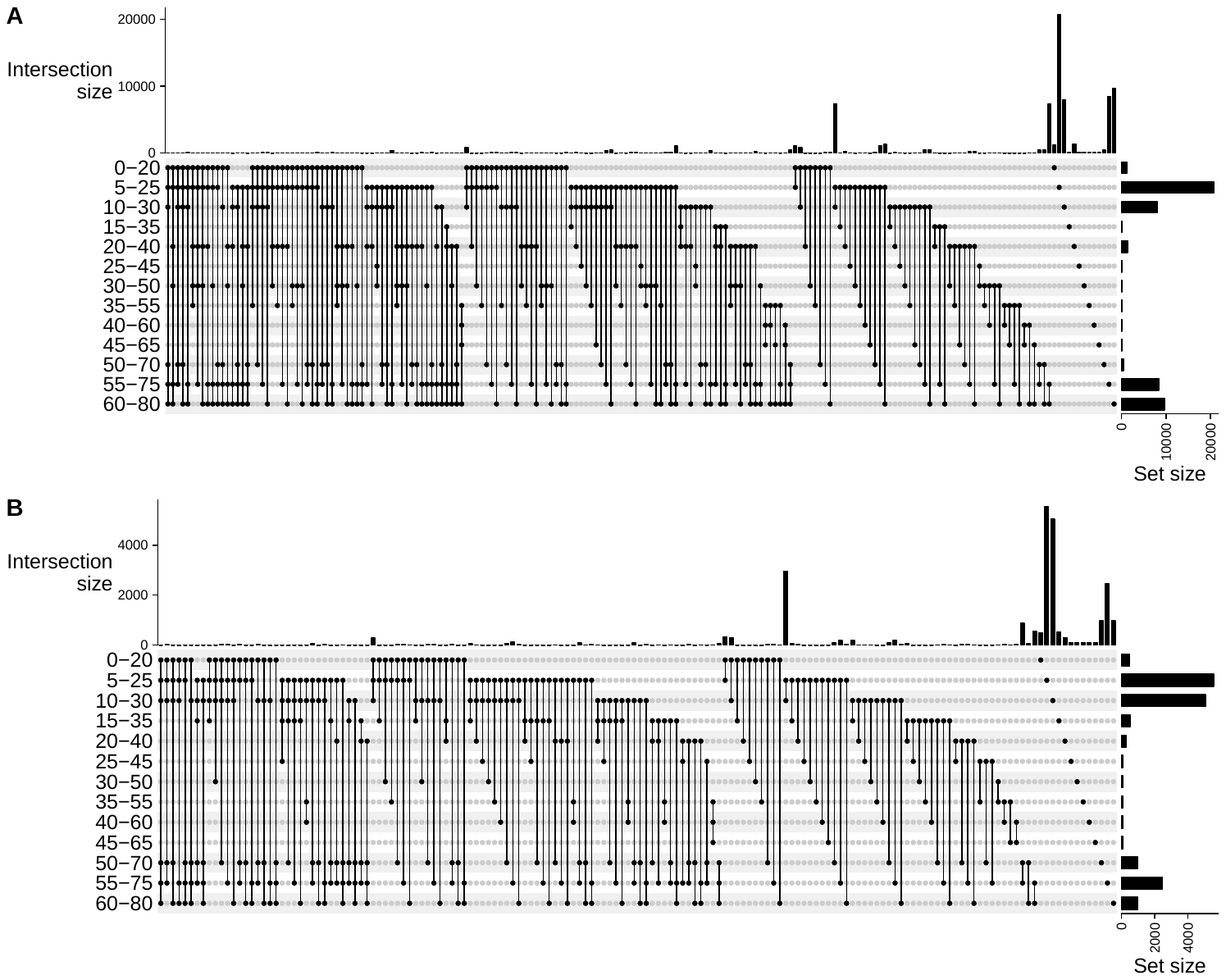

**Supplementary Figure 5.**

Epigenetic age calculated for all samples with the 4 epigenetic clocks used in this study. The other 5 clocks were not included because: Wu and McEwen’s clock can only be applied to young population; Belsky’s clock (DunedinPACE) only outputs epigenetic age acceleration; Teschendorff’s clock only calculates mitotic divisions; and Lu’s clock calculates telomere length. A) Hannum’s clock. B) Horvath’s clock. C) Levine’s clock. D) Shireby’s clock

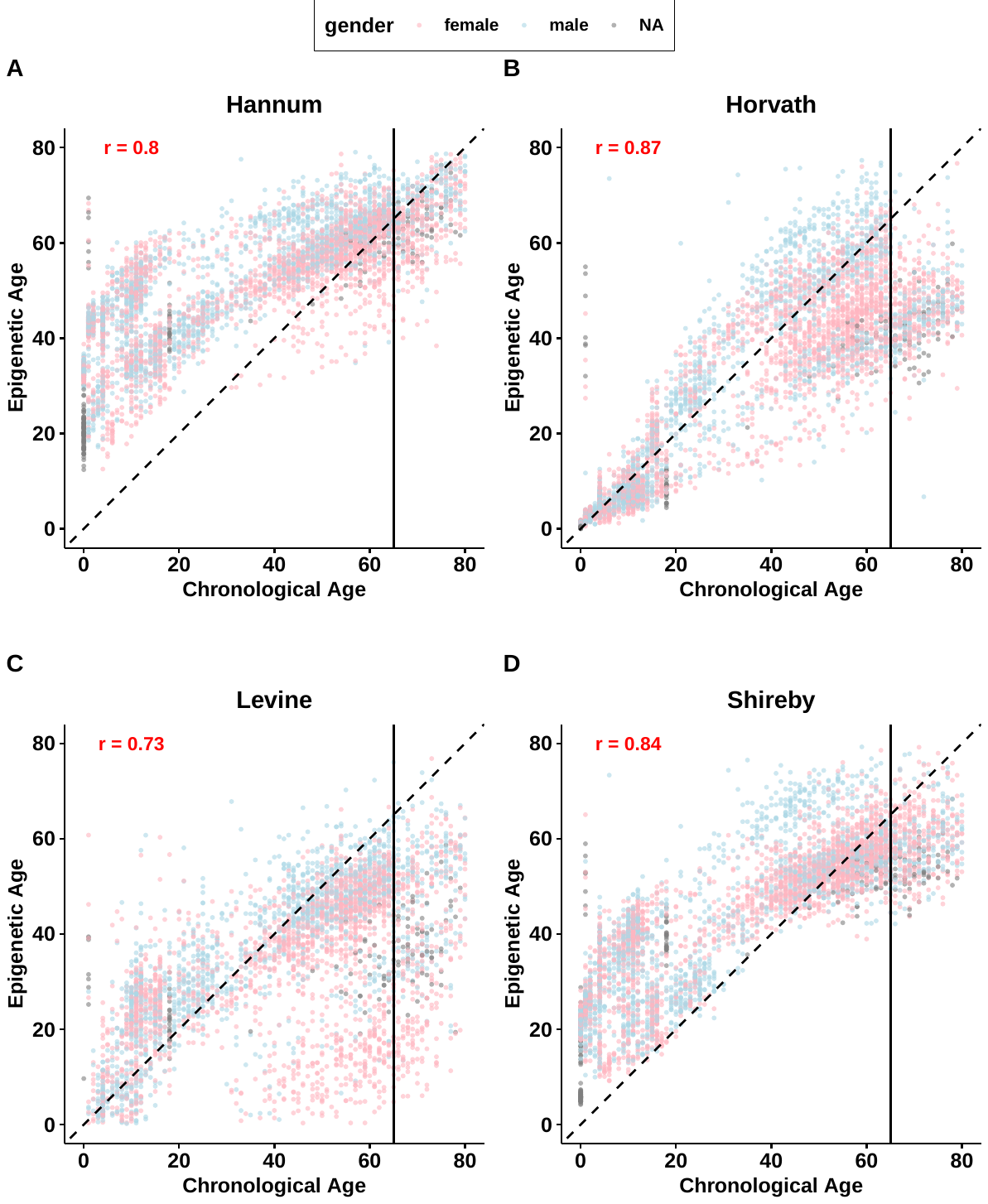

**Supplementary Tables**

**Supplementary Table 1**

Demographic characteristics of samples used in this study.

**Supplementary Table 2**

Summary of 23 datasets used in this study.

**Supplementary Table 3**

Summary of 9 epigenetic clocks used in this study.

**Supplementary Table 4**

Supplementary Table 4. Number of aaCpGs selected for whole cohort, male-only, female-only, and overlaps between male and female in different age windows prior to the addition of extra CpGs to ensure a minimum of 100 aaCpGs per window.
